# scContrast: A contrastive learning based approach for encoding single-cell gene expression data

**DOI:** 10.1101/2025.04.07.647292

**Authors:** Winston Li, Ghulam Murtaza, Ritambhara Singh

## Abstract

Single-cell RNA sequencing (scRNA-seq) captures gene expression at a individual cell resolution, which reveals critical insights into cellular diversity, disease processes, and developmental biology. However, a key challenge in scRNA-seq analysis is clustering similar cells across multiple batches, particularly when distinct sequencing protocols are used. In this work, we present scContrast, a semi-supervised contrastive learning method tailored for embedding scRNA-seq data from both plate- and droplet-based protocols into a universal representation space. By leveraging five simple augmentations, scContrast extracts biologically relevant signals from gene expression data while filtering out batch effects and technical artifacts. We trained scContrast on a subset of Tabula Muris tissues and evaluated its zero-shot performance on unseen tissues. Our results demonstrate that scContrast generalizes effectively to new tissues and outperforms the leading UCE approach in integrating scRNA-seq data from droplet- and plate-based sequencing protocols.

## 1 Introduction

Single-cell RNA sequencing (Jovic et al., 2022) (scRNA-seq) enables researchers to profile gene expression at the level of individual cells, permitting insights into cellular heterogeneity (Ximerakis et al., 2019), developmental processes (Qiu et al., 2022), and disease mechanisms (Mathys et al., 2019). A typical scRNA-seq analysis begins by clustering similar cells and identifying key marker genes for each cluster to map out differentiation pathways or uncover disease mechanisms (Su et al., 2022). However, clustering cells based on the full gene expression profile is particularly challenging due to the high dimensionality, sparsity, and batch effects inherent to scRNA-seq data (Bouland et al., 2023).

Existing computational methods for clustering scRNA-seq data can be broadly divided into three categories. The first category includes unsupervised approaches, which typically involve a highly variable gene selection followed by dimensionality reduction methods such as Principal Component Analysis (PCA) to get low-dimensional cell-level representations that cluster similar cells together. While such approaches work well for individual datasets, these approaches struggle to integrate data from multiple batches because they cannot account for batch effects or technical artifacts (Yuet al., 2024). To resolve these challenges, a second category of methods that includes supervised approaches such as scVI (Lopez et al., 2018) requires batch labels to assign similar representations to similar cells. However, these supervised approaches need to be retrained for every new integration, making them computationally expensive and impractical for large-scale datasets. More recently, inspired by the success of large foundational models such as ChatGPT (Brown et al., 2020), LLAMA (Touvron et al., 2023), and SAM (Kirillov et al., 2023), researchers have developed similar foundational models for scRNA-seq (Theodoris et al., 2023; Yang et al., 2022; Cui et al., 2024; Rosen et al., 2023). These foundational models are extremely large transformer-based architectures trained on atlas-scale datasets with a masked language modeling (MLM) objective. While these models promise to project arbitrary scRNA-seq data into a universal embedding space, it has been shown that these models struggle to effectively integrate data generated from different experimental protocols (Kedzierska et al., 2023).

This work introduces **scContrast**, summarized in Figure. 1, which is a semi-supervised contrastive learning approach designed to embed scRNA-seq data from different experimental protocols into a universal cell-level representation space. scContrast leverages a simple yet effective set of five augmentation functions, which simulate the technical artifacts, protocol biases, and batch effects characteristic of scRNA-seq assays. These augmentation functions guide scContrast to extract biologically meaningful features of single-cell gene expression profiles.

**Figure 1:**
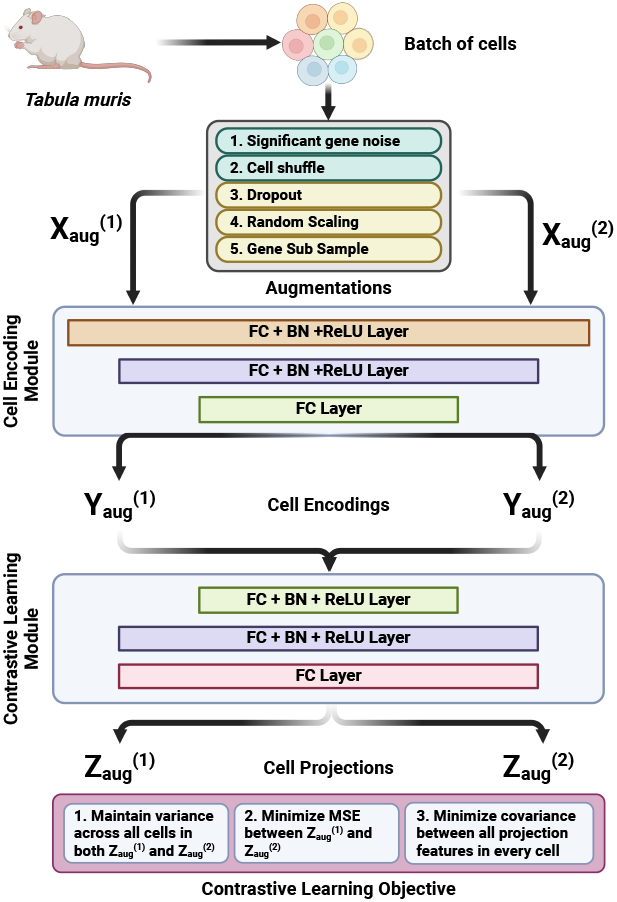
Overview of scContrast pipeline. First, given a batch of samples as a gene expression matrix, two augmented views are generated. Then, the views are encoded by a series of fully connected layers. Next, the encodings are projected into a higher dimension space by another series of fully connected layers. Finally, the projections are used in a contrastive learning objective that simultaneously maintain variance across samples, minimize MSE between augmented views, and minimize covariance between projection features.

**Figure 2:**
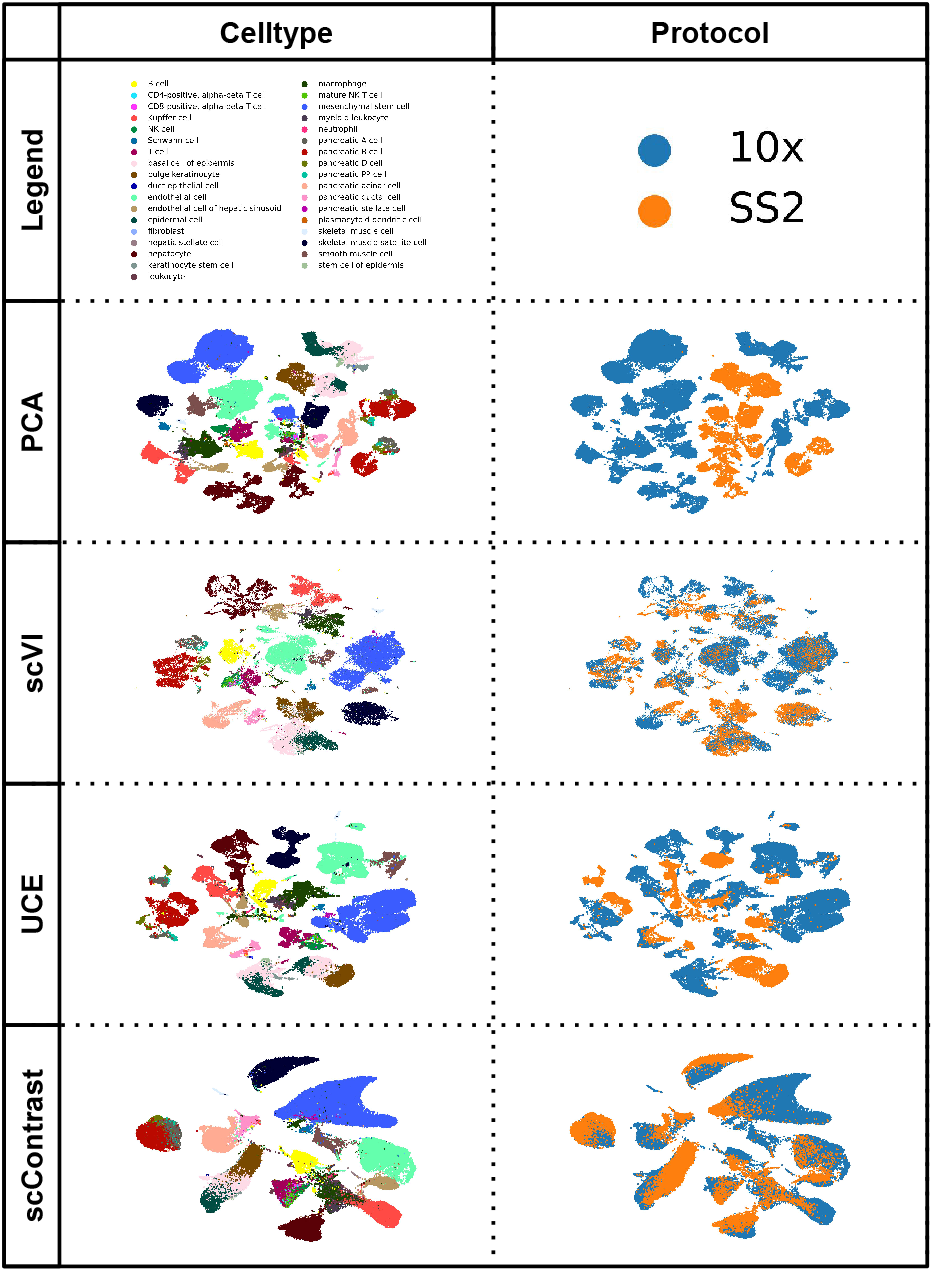
Qualitative comparisons of different models integrating scRNA-seq samples from plate (SS2)- and droplet (10x)-based sequencing platforms in unseen tissues in Tabula Muris

We trained scContrast on 21 out of 25 tissue samples from the Tabula Muris dataset (Consortium, 2018), which includes data from multiple donor mice sequenced using both droplet- and plate-based sequencing platforms. To simulate a true zero-shot setting, we tested scContrast on the remaining four held-out tissues. Through our evaluations, we found that scContrast generalizes well to unseen tissue samples and outperforms the state-of-the-art UCE model in integrating batches from different sequencing protocols. Specifically, scContrast scores 0.18 and 0.73 on the kBET and PCR batch integration metrics, while UCE scores 0.02 and 0.39, respectively. Thus, our findings highlight scContrast’s superior zero-shot cross-protocol batch integration capabilities. However, scContrast performs slightly worse on KMeans ARI and ASW bio-conservation metrics compared to UCE, suggesting opportunities for improvement in the future.

## 2 Method

As shown in Figure 1, scContrast has three main components:

### 2.1 Augmentations Module

The augmentations module take in a batch of cells from the Tabula Muris dataset to produce two augmented views 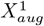 and 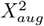 in a semi-supervised fashion. For each sample, the corresponding augmented views form a positive pair. This module relies on five augmentation functions that aim to perturb the gene expression data in a manner that preserves each sample’s cellular identity. Such functions can be sub-divided into two categories. The first category of augmentation functions require cell-type annotations (highlighted in green in Figure. 1). By leveraging the cell-type annotations of each sample, they ensure samples from the same cell-type maintain similar representations while challenging the model to distinguish between distinct cell-types. The significant gene noise augmentation injects noise into each cell’s most differentially expressed genes, which promotes robustness to expression variability in cell-type defining genes. The cell-shuffle augmentation permutes the gene expression profiles of samples of the same cell-type, which forces the model to treat similarly cells sharing an annotation.

The second category of augmentation functions does not rely on any annotations (highlighted with yellow color in the Figure. 1). These augmentation functions randomly dropout, subsample, or rescale gene expression across all cells to simulate random dropout events and sequencing protocol biases, consequently guiding the model to ignore these biases.

### 2.2 Cell Encoding Module

The core purpose of the Cell Encoding Module is to encode cells into a low-dimensional representation space that preserves biological semantic properties such as cell-type similarities, relationships, and developmental trajectories. Unlike existing scRNA-seq foundational models such as UCE (Rosen et al., 2023), Geneformer (Theodoris et al., 2023), scGPT (Cui et al., 2024), and scBERT (Yang et al., 2022) that have transformer blocks with self-attention, we rely on simple, fully connected neural networks to extract relationships between all genes in the cell. Specifically, we have two fully connected layers with batch normalization to stabilize the learning process and ReLU to capture non-linear relationships, followed by a single fully connected layer to produce cell encodings 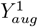 and 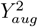. The entire Cell Encoding Module has 5.2 million parameters, which is 10—100 times smaller than existing scRNA-seq foundational models (Theodoris et al., 2023; Rosen et al., 2023).

### 2.3 Contrastive Learning Module

The Contrastive Learning Module is responsible for comparing the two augmented encodings generated by the Cell Encoding Module. It is inspired by VICReg (Bardes et al., 2022) and aims to ensure **V**ariance, **I**nvariance, and **C**ovariance among each batch of projected sample encodings to further enrich latent space representation. We use the same encoder and projection head for each augmented view.

More specifically, outputs from the Cell Encoding Module are first projected into 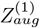 and 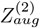, respectively, using a lightweight, trainable projection head. Then, a loss comprising three terms are calculated from these projected augmentations as shown in the equation below, where *X* and *Y* are 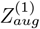 and 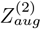, respectively:

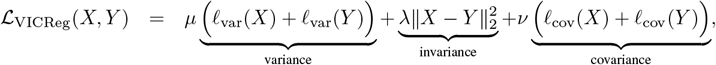

where the variance term is 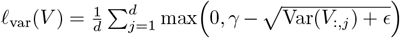, the invariance term is 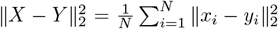, and the covariance term is 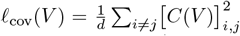, such that *V* ∈ {*X, Y*}, *γ* is the target standard deviation, *ϵ* is a small constant (e.g., 10^*−*4^) for numerical stability, and *C*(*V*) is the empirical covariance matrix of embeddings *V*, which is squared and summed over all off-diagonal elements. Coefficients *µ, λ*, and *ν* are tunable hyperparameters that weigh each loss term.

### 2.4 Implementation details

We implemented the entire pipeline in `Python==3.9.16` using `Pytorch==2.5.1+cu118` and `Pytorch-Lightning==2.4.0`. scContrast produces cell encodings with 128 dimensions and cell projections with 256 dimensions. We use the cell encodings *Y* for all of our evaluations and visualizations. We use cell projections 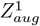 and 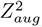 to calculate the VICReg (Bardes et al., 2022) contrastive loss with a batch size of 8192. We use AdamW optimizer with default parameters to calculate gradients used to update all the weights.

## 3 Experimental Setup

### 3.1 Datasets

We obtained scRNA-seq datasets from the Tabula Muris dataset, which is a comprehensive collection of single-cell transcriptomics data generated from both droplet-based (10x Genomics (Zheng et al., 2017)) and plate-based (Smart-Seq2 (Picelli et al., 2014)) sequencing platforms for *Mus musculus* (mouse). The Tabula Muris dataset contains 25 distinct tissues and has more than 350, 000 cells. In order to evaluate the performance of scContrast on out-of-distribution data in a zero-shot setting, we randomly selected four of the twenty-five tissues for testing and kept the rest for training scContrast.

We preprocessed all datasets by first filtering out all cells with ‘UKWN’/NULL annotations. Next, we normalized gene expressions of each cell by scaling the total counts to 10000. Finally, we applied a log-transformation to minimize technical artifacts (Su et al., 2022).

### 3.2 Baselines

In order to comprehensively compare the performance of scContrast against existing approaches, we include:

a. **PCA**: A fully unsupervised method that decomposes the cell-by-gene matrix into its principal components, producing low-dimensional cell representations.
b. **scVI** (Lopez et al., 2018): A supervised batch-integration approach that explicitly requires batch annotations. We train scVI with `n layers=3` and `n latent=128` to match the size and shape of the Cell Encoding Module of scContrast.
c. **UCE** (Rosen et al., 2023): A scRNA-seq foundational model with over 650 million parameters trained on an expansive corpus of scRNA-seq dataset containing over 36 million cells and spanning across eight diverse species. UCE can reportedly integrate scRNA-seq datasets with batch effects across unseen tissues and species without any finetuning. We use UCE with its default configurations without any finetuning to generate cell-level representations.

### 3.3 Evaluation Metrics

We evaluate the quality of cell encodings across bio-conservation and batch integration metrics defined by the `scib-metrics` package (Luecken et al., 2021), which is commonly used for bench-marking scRNA-seq models. For the sake of conciseness, we show two out of five metrics for each category and the rest of the scores in the Appendix (Table. 2, Table. 3).

**Table 1:** Quantitative comparison of different models on bio-conservation and batch integration of unseen tissues from Tabula Muris. The best score is bolded and the second best scores are underlined.

| Model | Bio-Conservation |  | Batch Integration |  |
| --- | --- | --- | --- | --- |
|  | KMeans ARI | ASW | kBET | PCR |
| PCA | 0.42 | <u>0.60</u> | 0.00 | 0.00 |
| scVI | 0.40 | <u>0.56</u> | **0.25** | **0.93** |
| UCE | **0.51** | **0.65** | 0.02 | 0.39 |
| scContrast | <u>0.43</u> | <u>0.60</u> | <u>0.18</u> | <u>0.73</u> |

**Table 2:** Additional quantitative comparison of different models on bio-conservation metrics of representation spaces of cells from unseen tissues from Tabula Muris. The best scores are bolded and the second best scores are underlined.

| Model | Bio-Conservation |  |  |  |  |
| --- | --- | --- | --- | --- | --- |
|  | Isolated Labels | KMeans NMI | KMeans ARI | ASW Labels | cLISI kNN |
| PCA | **0.67** | <u>0.76</u> | 0.42 | <u>0.60</u> | 1.00 |
| scVI | **0.67** | <u>0.76</u> | 0.40 | <u>0.56</u> | 1.00 |
| UCE | **0.67** | **0.80** | **0.51** | **0.65** | 1.00 |
| scContrast | <u>0.62</u> | 0.75 | <u>0.43</u> | <u>0.60</u> | 1.00 |

**Table 3:** Additional quantitative comparison of different models on batch integration metrics of representation spaces of cells from unseen tissues from Tabula Muris. The best score is bolded and the second best scores are underlined.

| Model | Batch Integration |  |  |  |  |
| --- | --- | --- | --- | --- | --- |
|  | ASW Batch | iLISI kNN | kBET | Graph Connectivity | PCR |
| PCA | 0.54 | 0.00 | 0.00 | 0.76 | 0.00 |
| scVI | **0.85** | **0.02** | **0.25** | **0.92** | **0.93** |
| UCE | 0.73 | 0.00 | 0.02 | 0.79 | 0.39 |
| scContrast | <u>0.77</u> | <u>0.01</u> | <u>0.18</u> | <u>0.80</u> | <u>0.73</u> |

We present the following four metrics:

a. **Bio-conservation metric ARI**: Calculates the Adjusted Rand Index from the KMeans-clustered embedding space of each model.
b. **Bio-conservation metric ASW**: Calculates the average silhouette width with respect to cell-type.
c. **Batch integration metric kBET**: Conducts the *k*-nearest-neighbor batch effect test, which rewards neighborhoods of embeddings having the same distribution of batch labels as that of the full dataset.
d. **Batch integration metric PCR**: Performs a principal component regression comparison by comparing the explained variance before and after integration.

## 4 Results

### 4.1 SCContrast successfully integrates gene expression data from plate- and droplet-based sequencing platforms in unseen tissues

scContrast is distinguished from existing approaches by its comparatively light model size, small training set, lack of explicit batch labels during training, and most importantly, its ability to generalize to unseen tissue samples in a zero-shot manner. For baselines, we choose PCA, scVI, and UCE to represent existing unsupervised, fully-supervised, and large-scale foundational model paradigms that are currently used for batch integration, respectively. For evaluation, we use four held-out tissues from Tabula Muris: skin, liver, pancreas, and limb muscle. These tissues feature diverse cellular landscapes ranging from organ-specific cells to common immune cells such (e.g. macrophages and resident variants). In order to visualize the quality of the cell representation spaces, we use UMAP (McInnes et al., 2020), which is a commonly employed visualization method. Moreover, we quantified these representations using ARI, ASW, kBET, and PCR.

As shown in Figure. 4.1, we find that both PCA and UCE struggle to integrate scRNA-seq data from droplet- and plate-based sequencing protocols. Specifically, from the UMAP visualizations, we observe that cells of the same cell-type but different sequencing protocol form very distinct and dense clusters in PCA; those of UCE tend be closer to each other, but they still do not integrate to form homogeneous clusters. In contrast, both scVI and scContrast successfully integrate across sequencing protocols: the plate-based SS2-colored embeddings (orange) overlay atop the droplet-based 10X-colored embeddings (blue), highlighting that representations of the same cell-type across both protocols homogeneously cluster together.

Quantitatively, UCE performs best on the bio-conservation metrics, highlighting its capacity to cluster cells of the same type closer to each other in the cellular representation space. However, the batch integration metrics reflect findings in Figure. 4.1, showing that both PCA and UCE perform worse than scContrast and scVI in integrating samples from different sequencing protocols.

While these results show that our contrastive learning-based approach is promising in learning a universal cell representation space, there is potential to further refine our approach to learn better representation spaces.

## 5 Discussion and Conclusion

In this work, we have demonstrated a semi-supervised contrastive learning-based approach scContrast for encoding single-cell RNA-sequencing data into a universal cell representation space. We compared our model against PCA, scVI, and UCE, which exemplify current unsupervised, fully supervised, and large foundational model approaches. Results demonstrate that PCA understandably struggles to integrate the batches, since protocol-inherent gene expression biases likely produce spurious axes of variability, which is exacerbated by PCA’s reliance on covariances. More interestingly, UCE also struggles, which may reflect the limitations of a purely masked modeling approach. Finally, while scVI performs comparatively best at cross-protocol integration, it explicitly requires batch labels, which scContrast does not.

In the future, we aim to extend scContrast across three dimensions. First, we aim to develop better augmentation functions to closer simulate batch effects and technical artifacts to improve scContrast’s zero-shot capabilities. Second, we plan to scale up our training datasets and explore deeper architectures that can learn richer cell representations. Lastly, we plan to investigate techniques like 𝕏-sample Contrastive Learning (Sobal et al., 2024) to impose hierarchical cell ontologies in the representation space.

## A Appendix

Supplementary metrics from `scib-metrics` are given in table 2 and table 3. Our scContrast model’s architecture is given here here.

~~~
scRNASeqE_VICRegExpander(
  (encoder): scRNASeqEncoder(
     (encoder): Sequential(
       (0): Linear(in_features=20116, out_features=256, bias=True)
       (1): BatchNorm1d(256, eps=1e-05, momentum=0.1, affine=True,
                        track_running_stats=True)
       (2): ReLU()
       (3): Linear(in_features=256, out_features=128, bias=True)
       (4): BatchNorm1d(128, eps=1e-05, momentum=0.1, affine=True,
                        track_running_stats=True)
       (5): ReLU()
       (6): Linear(in_features=128, out_features=128, bias=True)
     )
  )
  (projector): scRNASeqProjectionHeadExpander(
     (projectionHead): Sequential(
       (0): Linear(in_features=128, out_features=256, bias=True)
       (1): BatchNorm1d(256, eps=1e-05, momentum=0.1, affine=True,
                        track_running_stats=True)
       (2): ReLU()
       (3): Linear(in_features=256, out_features=256, bias=True)
       (4): BatchNorm1d(256, eps=1e-05, momentum=0.1, affine=True,
                        track_running_stats=True)
       (5): ReLU()
       (6): Linear(in_features=256, out_features=256, bias=True)
     )
  )
  (vicreg_loss_fn): VICRegLoss()
)
~~~

